# The plant-specific DDR factor SOG1 increases chromatin mobility in response to DNA damage

**DOI:** 10.1101/2021.11.03.466744

**Authors:** Anis Meschichi, Svenja Reeck, Adrien Sicard, Frédéric Pontvianne, Stefanie Rosa

## Abstract

Homologous recombination (HR) is a conservative DNA repair pathway in which intact homologous sequences are used as a template for repair. How the homology search happens in the crowded space of the cell nucleus is, however, still poorly understood. Here, we measured global chromosome and double-strand break (DSB) site mobility in *Arabidopsis thaliana*, using *lacO*/LacI lines and two GFP-tagged HR reporters. We observed an increase in global chromatin mobility upon the induction of DNA damage, specifically at the S/G2 phases of the cell cycle. DSB sites showed remarkably high mobility levels at the early HR stage, with a subsequent drastic decrease in mobility associated with the relocation of DSBs to the nucleus periphery. Importantly, the increase in mobility was lost in *sog1-1* mutant, a central transcription factor of the DNA damage response in plants. Our results indicate that repair mechanisms actively regulate chromatin mobility upon DNA damage, implying an important role for this process during the early steps of the DNA damage response.

## INTRODUCTION

Genome integrity is constantly threatened by internal and external stressors. Therefore, in response to DNA damage, eukaryotic evolved elaborate DNA-damage response (DDR) systems that comprise DNA-damage signaling processes and DNA repair^1^. Among the different types of DNA damage, double-strand breaks (DSBs) are particularly harmful for cells, leading potentially to chromosome rearrangements or loss of entire chromosome arms^2^. DSBs can be repaired by two main pathways, nonhomologous end joining (NHEJ) and homologous recombination (HR) (Jackson, 2002; West et al., 2004). NHEJ is achieved by stabilization and re-ligation of broken DNA ends, often with loss or mutation of bases. HR is a more complex and more conservative mechanism in which intact homologous sequences are used as a template for repair. HR most commonly occurs in S/G2 phases of the cell cycle in eukaryotic cells when sister chromatids are present, although homologous donor templates present elsewhere in the genome can also be used ^3–5^. Despite the vast knowledge about the molecular players involved in DNA repair via HR, the mechanisms behind the search and recognition of homologous sequences (“homology search”) is still not well understood. In yeast, large-scale movements of DSBs have been identified following DSB induction ^6–9^. Yet, the precise functions of these movements still remains poorly understood.

Plants are subject to particularly high levels of DNA damage resulting from dependence on sunlight for energy and exposure to environmental stresses (Rounds and Larsen, 2008). Moreover, plant development is mostly postembryonic with a late germline differentiation. It is, therefore, particularly interesting to understand the mechanisms that allow these organisms to cope with the constant assaults to their genome integrity. Indeed, plants have evolved a distinct DDR master regulator - SUPPRESSOR OF GAMMA RESPONSE 1 (SOG1). This transcription factor initiates a repair response by inducing genes involved in cell cycle arrest and repair, as well as in programmed stem-cell death in response to DNA damage^10–12^. While the molecular processes involved in DDR pathway have been extensively characterized also in plants, little has been done to address how chromatin mobility changes in response to DNA damage and in particular to DSBs. Here, we have used locus tagging systems and HR reporter lines to study chromatin mobility upon genotoxic stress with the DSB-inducer agent zeocin. We observed that in the presence of DSBs, both damaged and potentially undamaged loci increase the volume that they explore within the nuclear space. We showed that this increase in chromatin mobility occurs specifically during the G2 phase of the cell cycle and depends on the plant-specific DDR master regulator SOG1, implying an important role for chromatin mobility during the early steps of the DNA damage response.

## RESULTS AND DISCUSSION

To measure chromatin mobility in plant cells we used the *lacO*/LacI-GFP locus-tagging system ^13,14^ (Fig. 1A) and quantified foci mobility using a mean square displacement (MSD) analysis. This analysis robustly measures the mobility of diffusing, fluorescently tagged chromosomal loci and provides kinetic parameters describing loci motion^15,16^. We first tested our setup by measuring “steady-state” chromatin mobility levels for cells in the division versus differentiation zones of the *Arabidopsis thaliana* (*Arabidopsis*) root (Fig. 1B). Measurements of histone exchange dynamics had previously shown that cells at the division zone have a more dynamic chromatin state as compared to differentiated cells^17,18^. Consistently, we observed that chromatin mobility is also higher in cells from the division zone compared to cells from the differentiation zone (Fig. 1C). The radius of constrain (Rc), which indicates the nuclear volume within which a fluorescent spot can move, was also significantly higher in cells from the division zone (Fig. 1C). These results confirmed that our setup is suitable to unpick differences in chromatin mobility between cells. In *Arabidopsis* root, differences in nucleus size are often evident, not only between nuclei from the division and differentiated zones but also within the meristem itself. As such, we thought to verify if our MSD measurements would be affected by differences in nucleus size. Within the meristematic region from the root, cells have the same ploidy level (diploid), but nuclei of atrichoblast cells are considerably bigger than that of trichoblast cells (Fig. 1D). Nevertheless, these two cell types show the same chromatin mobility and radius of constraint (Fig. 1E), ruling out that the nuclear volume *per se* could affect overall chromatin mobility levels.

**Figure 1:**
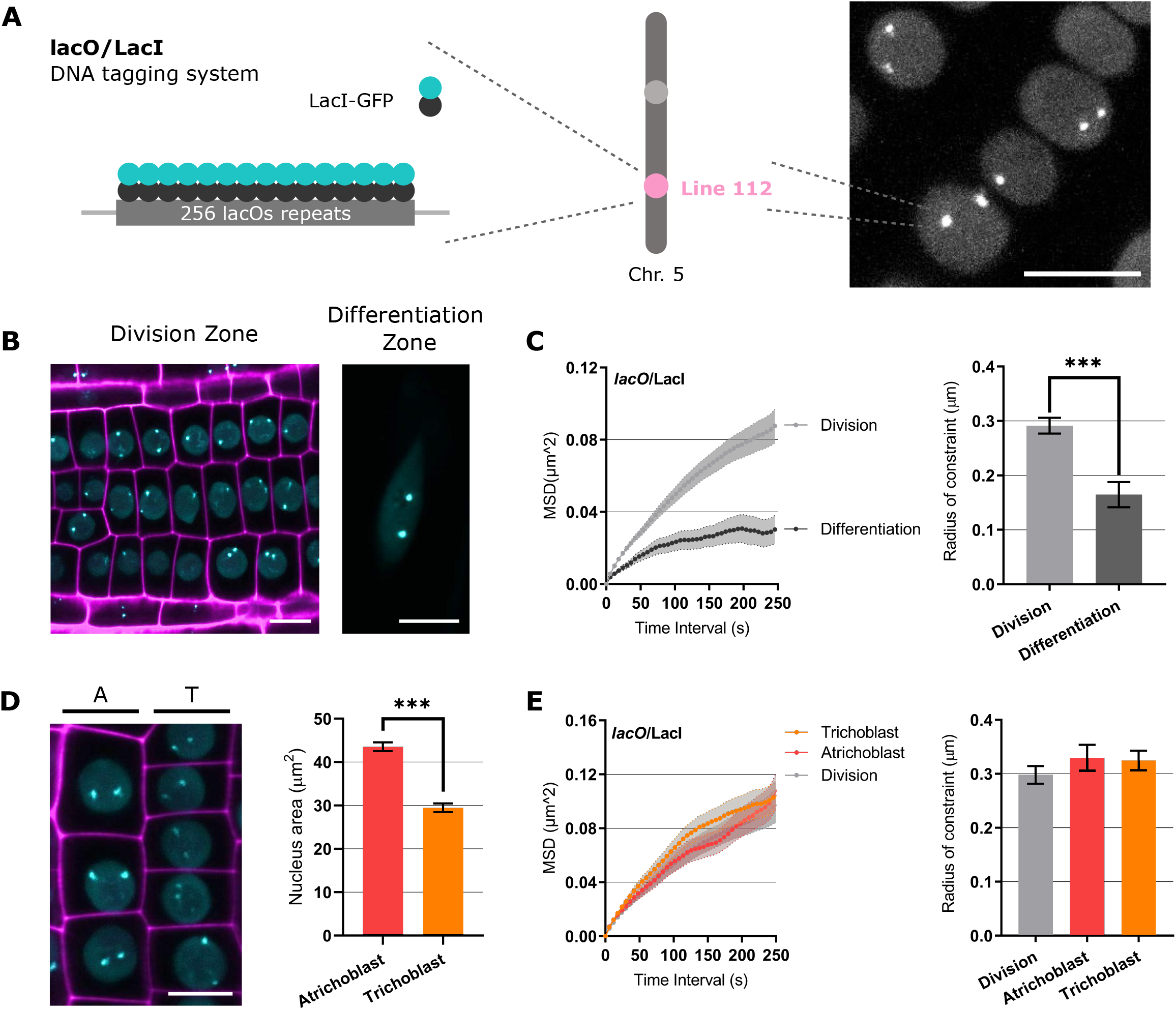
MSD analysis of *lacO* foci in different cell types in Arabidopsis thaliana root. (A) Schematic representation of the lacO/LacI system. A *lacO* repeat array was integrated into chromosome 5 (line112) and detected by expression of the LacI protein fused to GFP. The image on the right corresponds to z-projected images from root epidermal nuclei expressing the referred construct. Scale bar, 10μm (B) Representative images of the Arabidopsis root epidermal cells in division (left image) and differentiation zone (right image) showing nuclear signal with *lacO*/LacI foci (cyan). Propidium Iodide (PI) staining (magenta). Scale bar, 10μm. (C) MSD analysis of *lacO*/LacI lines based on time-lapse experiments of nuclei in the division (n=116 nuclei) and differentiated zone (n=21 nuclei). 3D stacks were taken at 6sec intervals for 5min. The radius of constraint was calculated from MSD curves. Values represent means ± SEM. Student t-test, ***P < 0.001. (D) Left: Representative images of atrichoblast (A) and trichoblast (T) in the division zone showing nuclear signal with *lacO*/LacI foci (cyan). Propidium Iodide (PI) staining (magenta). Scale bar, 10μm. Right: Histogram of nuclear areas (μm2) from atrichoblast and trichoblast cells. Atrichoblast (n = 53 nuclei); red, Trichoblast (n = 57 nuclei); orange. (E) Left: MSD analysis of *lacO*/LacI lines based on time-lapse experiments of nuclei in the atrichoblast (n=36 nuclei) and trichoblast (n=61 nuclei). Right: Radius of constraint calculated from MSD curves. Values represent means ± SEM. Student t-test, ***P < 0.001.

Because HR requires pairing of the broken DNA molecule with a homologous intact template, we tested whether *Arabidopsis* cells actively regulate the chromatin mobility in response to DSBs to promote conservative repair mechanisms. We induced DNA damage by incubating 6d-old seedlings with the DSB inducer zeocin for 24h (Fig. 2A). This treatment led to the upregulation of the DDR responsive genes PARP2, RAD51 and BRCA1, indicating that the HR was effectively stimulated (Supplementary Fig. 1A). This provided us with a system to induce different levels of DNA damage and repair mechanisms. We further focused our analysis on cells within the division zone since previous studies showed that the principal actors of HR, RAD51 and RAD54 are mainly expressed in these cells^19,20^. MSD analysis revealed that *lacO*/LacI foci mobility was not changed upon low concentrations or shorter times of zeocin incubation but increased significantly with high concentrations of zeocin for 24h (Fig. 2B, Supplementary Fig. 2A B). Importantly, the effect seen at the higher concentration was not due to DNA damage-induced programmed cell death as tested by PI staining (Supplementary Fig. 3). Only stem cells and their early descendants, which are known to be highly sensitive to DNA damage^21^, showed PI-positive cells but not the epidermal cells used in our chromatin mobility analysis. We also tested other DSB inducer chemicals, namely mitomycin C (MMC). A similar increase in chromatin mobility was observed in response to MMC treatment (Supplementary Fig. 4), showing that this is a general response to DSB induction.

**Figure 2:**
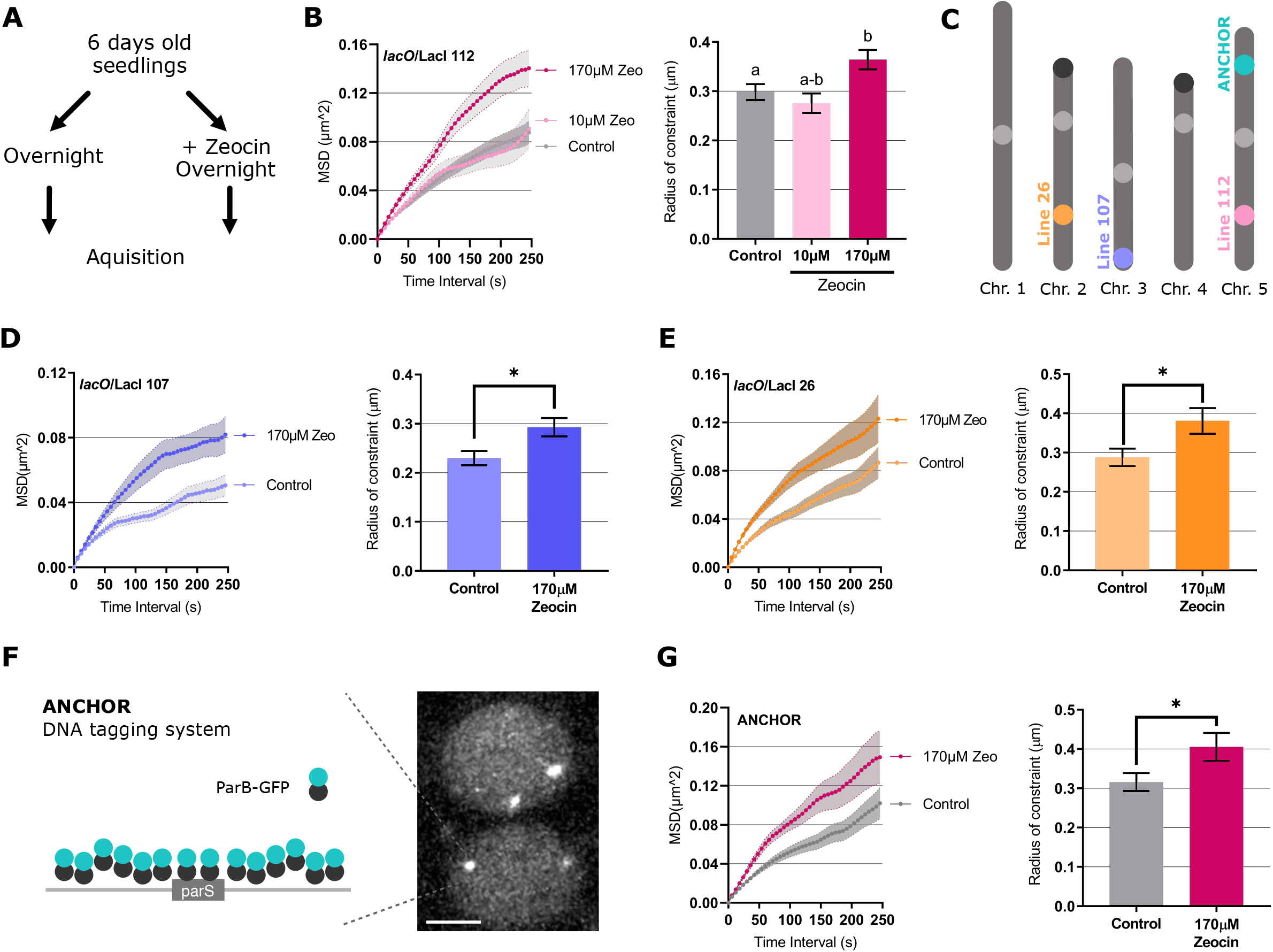
Chromatin mobility increases upon high DNA damage levels. (A) Scheme illustrating the experimental setup. (B) MSD analysis of *lacO*/LacI line 112 based on time-lapse experiments of nuclei in different zeocin concentrations. 10μM (n=97 nuclei); 170μM (n=93 nuclei). Radius of constraint were calculated from MSD curves. Values represent means ± SEM. Letters indicate one-way ANOVA followed by Bonferroni’s correction (p < 0.05). (C) Chromosomal positions of *lacO*/LacI lines as reported previously 9,10. Line 26 and line 107 are respectively inserted in chromosomes 2 and 3. The ANCHOR construct is inserted in chromosome 5. The NORs are marked as black circles and the centromeres as light grey circles. (D) MSD analysis of *lacO/*LacI line 107 based on time-lapse experiments of nuclei in control conditions and zeocin treated plants with 170μM. Control (n=53 nuclei), 170μM (n=48 nuclei). Bottom: Radius of constraint calculated from MSD curves. Student t-test, *P < 0.05. (E) MSD analysis of *lacO*/LacI line 26 based on time-lapse experiments of nuclei upon zeocin. Control (n=52 nuclei), 170μM (n=52 nuclei). Radius of constraint calculated from MSD curves. Student t-test, *P < 0.05. (F) Left: Schematic representation of the ANCHOR system. *parS*-ParB:GFP interactions and oligomerization along the flanking genomic region. ParB-GFP can directly bind to *parS* sequence as a dimer and along the flanking genomic region. Right: Representative image of epidermis nuclei in the division zone. Scale bar, 5μm. (G) MSD analysis of ANCHOR line based on time-lapse experiments of nuclei upon zeocin treatment. Control (n=54 nuclei), 170μM (n=22 nuclei). Radii of constraint calculated from MSD curves. Student t-test, *P < 0.05.

In order to verify if the increase in chromatin mobility observed upon zeocin treatment was specific for the particular *lacO* insertion site (line112) or a response at the global chromatin level, we analyzed additional *lacO*/LacI lines with insertions at different chromosomal locations (Fig. 2C). In control conditions, line 26 shows the same chromatin mobility as line 112, whereas line 107 showed significantly lower chromatin mobility and Rc (Supplementary Fig. 5). The lower mobility in line 107 could be linked to the transgene insertion at the subtelomeric region which are known to physically interact at the nucleolar periphery in *Arabidopsis*^22–24^ (Fig. 2C). Upon treatment with high zeocin concentration, all lines showed a significant increase in chromatin mobility and Rc (Fig. 2D and E), indicating that chromatin mobility increases globally in the nucleus in response to DNA damage. We also tested whether these results could be an artefact of the *lacO*/LacI system itself. For that, we performed the same experiments using another locus tagging system - the ANCHOR system (ParB-*parS*)^25^ (Fig. 2F). The ANCHOR line showed a similar increase in chromatin mobility (Fig. 2G).

Existing evidence in several systems show that cell cycle arrest upon DNA damage is often used by cells to facilitate DNA repair before cell division^26–28^. Since DNA content and cohesion differ in different cell cycle phases, we sought to test if changes in cell cycle dynamics (i.e. the proportion of cells in different cell cycle phases) could explain the increased chromatin mobility observed in response to DNA damage. To test this hypothesis, we crossed the *lacO*/LacI (line 112) with the S/G2 reporter CDT1a::RFP^29^ (Fig. 3A, B). To first verify that this setup was working as expected, we quantified the ratio of cells in S/G2 in root epidermal cells treated with Hydroxyrea (HU), a drug known to block cells in S phase^30,31^. Indeed, we observed that there was a higher proportion of cells in S/G2 in HU samples (Fig. 3C). Consistent with previous studies^32^, treatment with 10μM zeocin significantly increased the number of cells in G2/S (Fig. 3C). However, with the highest concentration of zeocin (170μM), the ratio of cells in S/G2 phase decreased to half in comparison with control conditions (Fig. 3C), suggesting an accumulation of cells in G1. Thus, it became important to determine if G1 cells had different chromatin mobility compared to S/G2 cells. MSD analysis revealed that cells in the S/G2 phase (CDT1a-RFP positive cells) showed lower chromatin mobility than G1 cells (Fig. 3D). Similarly, HU-treated cells, showed lower chromatin mobility, most likely due to cells being arrested in the S/G2 phase (Fig. 3E). These results revealed that an accumulation of cells in G1, could potentially explain the increased mobility observed in response to DSBs. If this is the case, we hypothesized that we should not see differences when comparing cells at the same stage of the cell cycle with or without zeocin. We, therefore, measured the chromatin mobility specifically at G1 and S/G2, in control conditions and upon treatment with different concentrations of zeocin. We observed a significant increase in chromatin mobility in cells at S/G2 after zeocin treatment, whereas cells in G1 did not show any significant change (Fig. 3F and G). We concluded that the increased mobility observed in response to DNA damage at high zeocin concentrations (170μM) could be both a result of an accumulation of cells in G1 and a specific increase in chromatin mobility at S/G2 phase. This observation is consistent with the idea that HR is particularly relevant in G2 when sister chromatids have been synthesized and suggests that increased chromatin mobility may be important during this stage.

**Figure 3:**
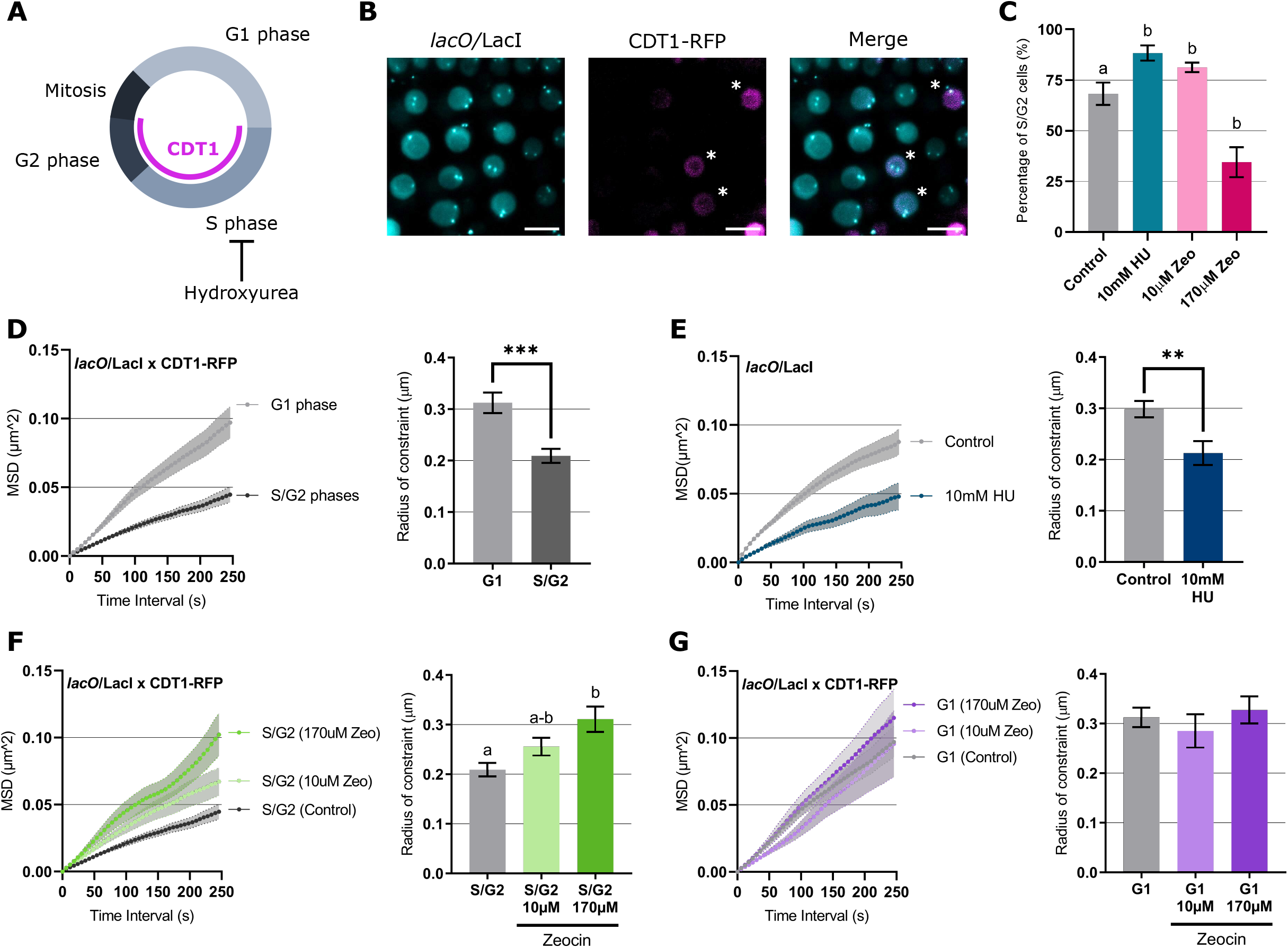
Chromatin mobility increases specifically during in S/G2 phases in response to DNA damage. (A) Schematic representation of cell cycle progression with the CDT1-RFP signal displayed in cell in S/G2. (B) Representative images of nuclei from *lacO*/LacI line 112 crossed with CDT1-RFP, *lacO*/ LacI (cyan) CDT1-RFP (magenta). Stars represent cells in S-G2. Scale bar, 10μm. (C) Percentage of S/G2 cells per root in control conditions and upon 10μM hydroxyurea, 10μM and 170μM Zeocin. (d-h) MSD curves and corresponding Rc histograms for: (D) MSD measurements of nuclei in G1 (n=62 nuclei) and S/G2 phase (n=67 nuclei); (E) *lacO*/LacI lines based on time lapse experiments of nuclei upon 10μM HU treatment phase (n=28 nuclei); (F) S/G2 cells upon different zeocin concentration (10μM (n=60 nuclei); 170μM (n=49 nuclei)); (G) G1 cells upon different zeocin concentration (10μM (n=35 nuclei); 170μM (n=50 nuclei)); Values represent means ± SEM. Student t-test, *P < 0.05 **P < 0.01 ***P < 0.001. Letters indicate one-way ANOVA followed by Bonferroni’s correction(p < 0.05)

In yeast, as in plants, studies have shown that HR is executed mainly during S/G2 phases of the cell cycle^26,33^. Because the increase in mobility upon zeocin treatment was specific to S/G2, we decided to investigate the mobility of DSBs during HR. Homologous recombination is divided into two main phases: the presynaptic phase, which includes 5’-end resection and homology search, and the synaptic phase, which includes the strand invasion for homologous strand pairing (Fig. 4A)^34^. The two main actors of HR, RAD51 and RAD54, function respectively in the initiation of the strand invasion and at the strand exchange reaction that finalizes the repair^35^. We wanted to investigate how the increase in chromatin mobility is placed in relation to these two phases. By performing an eight-hour time course experiment on RAD51-GFP and RAD54-YFP lines after induction of damage with 10μM zeocin, we were able to visualize the appearance of foci with accumulations of these proteins in the nucleus. This revealed that RAD51-GFP foci were formed approximately 1h30min after DSB induction, whereas RAD54-YFP foci appeared later, at around 5h after treatment (Fig. 4B). From this experiment, we could confirm that RAD51 interacts first with DSBs, while RAD54 comes in later. To investigate the mobility of foci tagged with these proteins, we treated RAD51-GFP and RAD54-YFP plants with 10μM zeocin, (Fig. 4C). The MSD analysis revealed that only RAD51 showed significantly higher mobility than *lacO*/LacI foci (Fig.4D), showing that high mobility levels seem to happen at early HR stages. Previous studies have shown that RAD54 foci relocate to the nuclear periphery after γ-irradiation^32,33^. Therefore, our MSD results for RAD54 may correspond to a mixture of foci located at the nuclear periphery and non-periphery. To test if RAD54 at the different nuclear compartments behaved differently, we determined the MSD for RAD54 foci at these two nuclear locations (Fig.4E). The results showed that non-peripheric RAD54 foci have much higher mobility than the foci at the periphery (Fig.4F), revealing that RAD54 foci can, depending on their location, have mobilities similar to those of RAD51. Moreover, these results highlighted that large changes in chromatin mobility occur during the repair process – a strong increase in DSB mobility is observed in the early HR phase, with a subsequent drastic drop in mobility associated with the relocation of DSBs to the nuclear periphery. This relocation to the nucleus periphery has been associated with different possible roles - to bring homologous sequences together, thereby reducing the 3D search to a 2D scale ^36^; or due to the fact that the repair machinery may specifically interact with nucleopores ^37^.

**Figure 4:**
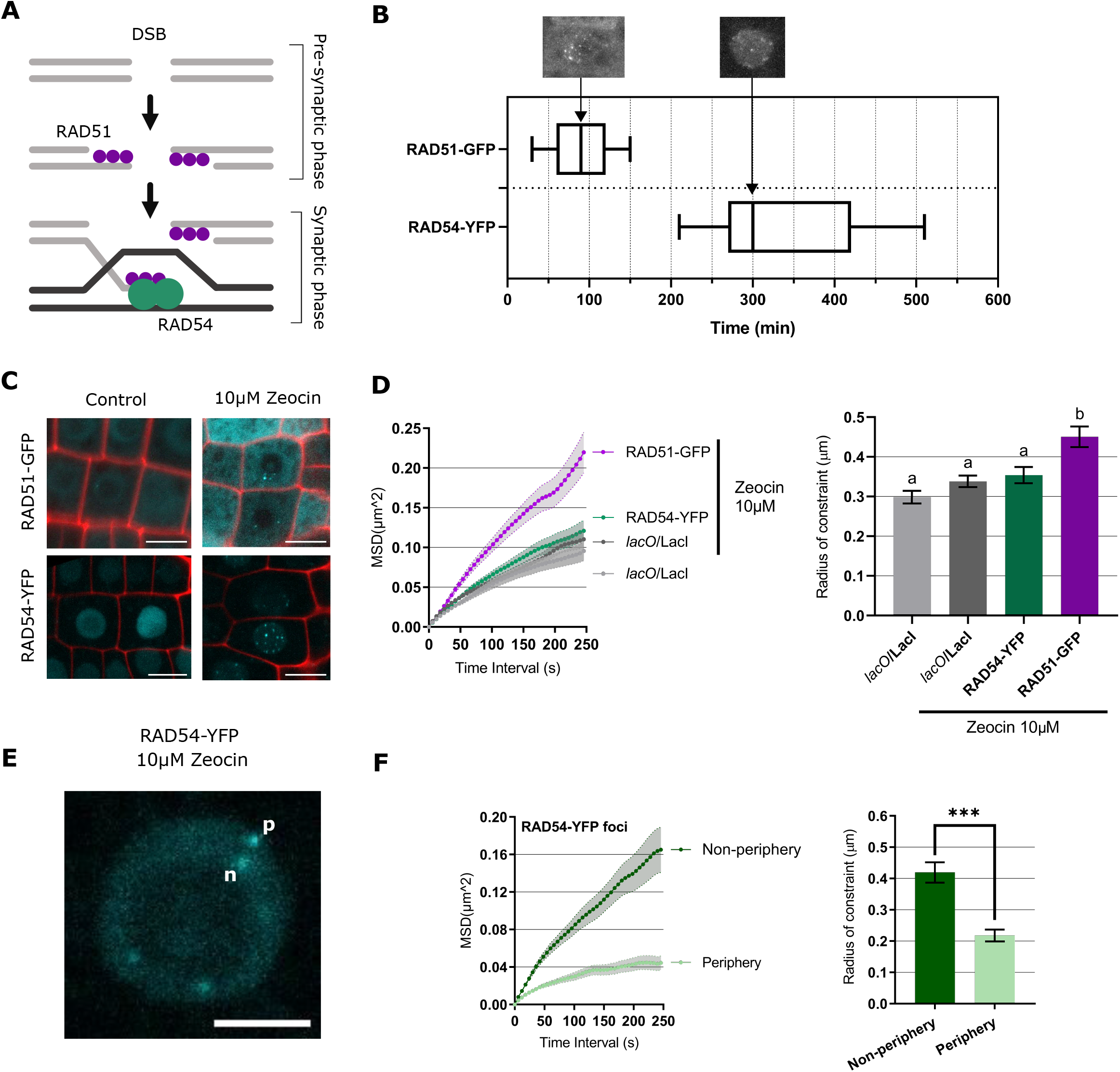
DSB mobility is higher at early HR. (A) Schematic representation of the critical steps of homologous recombination. Rad51 (purple) assemble onto the single-stranded DNA (ssDNA) formed after exonucleation of DNA double-strand break (DSB) ends to form a filament, which is known as the presynaptic filament. After searching for DNA homologous sequence, the presynaptic filament binds the DNA template to form the synaptic structure with RAD54. The ssDNA invades the homologous region in the duplex to form a DNA joint, known as the displacement (D)-loop promoted by Rad54 (green). (B) Time-lapse experiment of the formation of RAD51-GFP and RAD54-YFP foci in Arabidopsis nuclei, which was imaged every 30min after zeocin treatment. Timeline of RAD51 and RAD54 foci formation for 8h. Error bars indicate the standard error. At least four roots were counted for each group. (C) Representative images of root epidermal cells showing foci formation in RAD51-GFP and RAD54-YFP plants after 10μM zeocin treatment for 48h. Propidium Iodide (PI) staining (red). Scale bar, 10μm. (D) MSD analysis of RAD51 (n=64 nuclei) and RAD54 (n=64 nuclei) foci and *lacO/*LacI (line112) (n=109 nuclei) plants upon 10μM zeocin. Radius of constraint calculated from MSD curves. (E) Representative images of root epidermal nuclei with RAD54 foci located on the nuclear periphery (p) and non-periphery (n). Scale bar, 5μm (F) MSD analysis of RAD54 foci in the periphery (n=24 nuclei) and non-periphery (n=30 nuclei) upon 10μM zeocin. Radius of constraint calculated from MSD curves. Values represent means ± SEM. Student t-test, ***P < 0.001. Letters indicate one-way ANOVA followed by Bonferroni’s correction (p < 0.05).

Tracking chromatin movement, using DNA labelling tools and HR reporter lines, showed an increase in mobility upon DNA damage. Next, we wanted to determine whether the increase in mobility was actively regulated by the DDR pathway. For that we quantified *lacO*/LacI (line 112) mobility in *sog1-1* mutant, in which DDR is abolished. MSD analysis in *sog1-1* mutant revealed no increase in mobility upon treatment with high zeocin concentration, indicating that the increase of mobility seen in the WT (SOG1+/+ progeny from the F1) was dependent on SOG1 and thus on DDR activation (Fig. 5A, B). However, it is important to rule out that the lack of response to zeocin treatment was not due to a change in the cell cycle dynamics in this mutant. Indeed, in *sog1-1* the cell cycle arrest upon DNA damage is compromised ^32,38,39^ and a loss of G1-arrested cells could potentially explain the results observed. We used EdU staining to check if, under our zeocin treatment conditions, *sog1-1* cells were not being arrested in G1 (Fig. 5C-E; Supplementary Fig. 7). The results showed that also in *sog1-1* there is a substantial reduction in EdU staining upon zeocin treatment, indicating that cells are also being accumulated at G1 although to a less extent than in the WT. We therefore decided to further analyse the chromatin mobility in G1 and S/G2 in *sog1-1* mutant. Given the complexity of this line, with several T-DNA insertions, instead of crossing it with CDT1a::RFP reporter we used nuclear area as a proxy for cell cycle stage taking as a reference CDT1 labelling (Supplementary Fig. 8). This analysis revealed that in *sog1-1* at both G1 and S/G2 stages of the cell cycle there is no increase in mobility upon zeocin treatment (Fig. 5F-H). These results demonstrate that SOG1 is required for the increase in chromatin mobility induced by zeocin treatment indicating that this phenomenon is actively regulated during the early steps of the response to DNA damage and not a physical by-product from extensive DNA “fragmentation”.

**Figure 5:**
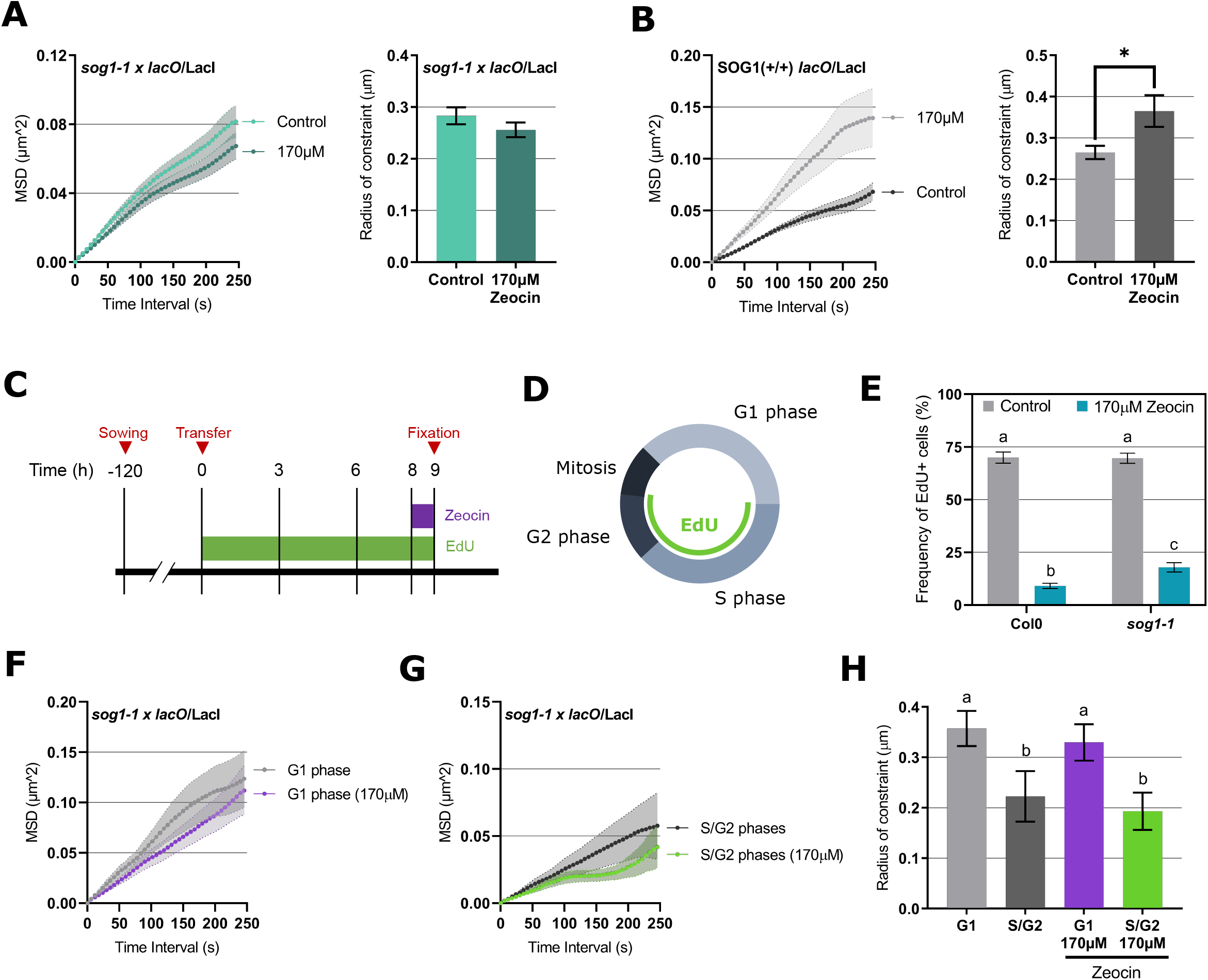
SOG1 is required for the increased chromatin mobility. (A) MSD analysis of *lacO/*LacI line 112 crossed with *sog1-1* (Control (n=83 nuclei); 170μM zeocin (n=91 nuclei)). (B) MSD analysis of SOG1+/+ *lacO/*LacI progeny from crossing with *sog1-1* (Control (n=59 nuclei); 170μM zeocin (n=29 nuclei)). Radius of constraint calculated from MSD curves. Student t-test, *P < 0.05. For all MSD curves, values represent means ± SEM. (C) Simplified schematic representations of the protocols corresponding to the EdU labelling experiment. (D) Schematic representation of cell cycle progression with the EdU signal displayed in cell in S/G2. (E) Proportion of EdU labelled cells in one root tip in Col-0 and *sog1-1* in control conditions and zeocin treated plants with 170μM. (F) MSD analysis of *lacO/*LacI line 112 crossed with *sog1-1* nuclei in G1 phase (Control (n=23 nuclei); 170μM zeocin (n=25 nuclei)). (G) MSD analysis of lacO/LacI line 112 crossed with *sog1-1* nuclei in S/G2 phase (Control (n=10 nuclei); 170μM zeocin (n=12 nuclei)). (H) Radius of constraint calculated from MSD curves. Values represent means ± SEM. Letters indicate one-way ANOVA followed by Bonferroni’s correction (p < 0.05).

Our analysis of chromatin movement has revealed that an increase in chromatin mobility occurs in response to DSBs in Arabidopsis. Similar responses have been observed in yeast and animal cells, pointing towards a general mechanism of response to DSBs across kingdoms^6,40–42^. The actual function of such increase in mobility has not been fully uncovered but some studies support the idea it may increase the probability of an encounter between the break and the repair template^43,44^. Despite the DNA repair machinery being highly conserved between eukaryotes, some of the important animal regulators, such as the tumor suppressor p53, have not been found in plants. Its function is instead served by the plant-specific DDR master regulator SOG1. Interestingly, we have been able to show that in plants the increase in chromatin mobility is dependent on SOG1 function. These results suggest that the increase in chromatin mobility, was conserved in evolution, as a response to DNA damage through the action of different molecular players. Further studies are now required to determine the mechanisms downstream of SOG1.

## MATERIAL AND METHODS

### Plant lines and growth conditions

Mutants and transgenic lines used in this study come from the following sources: *sog1-1*^10^, RAD51-GFP^19^, RAD54-eYFP^45^, Cytrap line^29^, *lacO*/LacI lines^13^, ANCHOR line^25^. All mutants and transgenic lines are in Columbia background.

To visualize S/G2 cells, *lacO*/LacI line 112 was crossed to Cytrap line, and the resulting F2 plants were selected on MS plates containing 50 mg/L of kanamycin (Sigma-Aldrich, catalogue number K1377). Because the G2/M-marker CYCB1;1 is strongly expressed during DNA damage^46^, the selected F2 were screened only for LacI-GFP and CDT1a-RFP.

Seeds were sterilized in 5% v/v sodium hypochlorite for 5 min and rinsed three times in sterile distilled water. Seeds were stratified at 4°C for 48 h in the darkness. Seeds were then plated on Murashige and Skoog (MS) medium and then grown in 16 hours light at 25°C in vertically oriented Petri dishes. The roots were observed after 6 to 7d of incubation, depending on the experiment.

### Genotoxic treatment

To induce DNA damage response, 5-to 6-day-old seedlings were transferred in MS medium without or with 100 μM mitomycin C (MMC); 2, 10, 50, 100 or 170 μM zeocin or 10 mM hydroxyurea (HU) and treated for 2h, 6h, or 24h. Each chemical was obtained respectively from Fisher Scientific (catalogue number 2980501), Invitrogen (catalogue number R25001) and Sigma-Aldrich (catalogue number H8627-1G).

### Microscopy

For root staining with propidium iodide (PI) 6 to 7-d-old seedlings were mounted in water between slide and coverslip and sealed with 0.12-mm-thick SecureSeal Adhesive tape (Grace Bio-Labs) to reduce drift drying during imaging.

For EdU staining, samples were imaged using a Zeiss LSM800 inverted microscope, with a 63x water-immersion objective (1.20 NA) and Microscopy Camera Axiocam 503 mono; Alexa499 were detected using a 488nm excitation filter by collecting the signal between 505-550 nm. For DAPI, an excitation filter 335-383nm was used, and the signal was detected between 420-470 nm

### Mean square displacement

For all MSD experiments, time-lapse imaging was performed every 6 s, taking a Z-stack of 3 μm spread through 1μm slices for 5 min, with a 512 × 512 pixels format with a 1-2× zoom factor. All images were analyzed using Fiji software (NIH, Bethesda, MD, http://rsb.info.nih.gov/ij/) (Sage et al. 2005) and with the plugin SpotTracker 2D (obtained from http://bigwww.epfl.ch/sage/soft/spottracker). Images were analyzed as described in Meschichi *et al*., 2021.

### Expression Analysis Using Real-Time RT-PCR (qPCR)

Seedlings grown for 7 d were harvested, and total RNA was extracted using the Trizol reagent (Invitrogen). A total of 1 μg of RNA was treated with TURBO DNase (Life Technologies) and used for cDNA synthesis (Superscript IV; Life Technologies). The resulting cDNA was diluted 10 times and used for quantitative PCR using a Bio-Rad iCycler Thermal Cycler iQ5 Multicolor Real-Time and HOT FIREPol® EvaGreen® qPCR Mix Plus (Solis Biodyne). For data normalization, the data were first normalized to the PP2A2 reference gene, and the values from two independent samples were normalized to the average Delta Ct value Col-0 level or control condition (2^-ΔΔCt Method). The final values presented are given as the mean ± SD from three independent samples. Minus RT (no reverse transcriptase control) controls were set up to make sure the values reflect the level of RNA and not DNA contamination. The standard Student’s t-test was used to determine the statistical significance of the results. The primers used are listed in Supplementary Tables 2.

### EdU labelling

Five-day-old seedlings were grown on solid medium containing 20 μM EdU, followed by a 1h incubation in EdU and 170uM zeocin. Roots were fixed in 4% paraformaldehyde (PFA) for 30 min, and washed three times with 1 × PBS. The roots were transferred to slide and covered by a glass cover slip, then squashed, and immediately deepen in liquid nitrogen for few seconds. The cover slips were removed and the roots were left to dry at room temperature for 30 min. The samples were washed with PBS + BSA (Bovine Serum Albumin) 3% (w/v), and incubated with a ClickIt Buffer (PBS 1X pH7.4, CuSO4 100mM, Ascorbate 1M, Alexia fluor azide, 2uM) solution in the dark for 15 minutes. Samples were washed once in 1X PBS + BSA 3%, followed by DAPI staining for 15 minutes in the dark. Samples were washed twice with PBS 1X pH 7.4 and mounted in vectashield (Vector Laboratories).

### Statistical analysis

For statistical analysis, we used the GraphPad Prism 8.3 software. Data set were tested for normality using the Shapiro–Wilk test. Statistical significance was determined by using the standard student t-test (two-tailed) and one-way ANOVA (multiple comparisons with Bonferroni correction. All experiments were performed in several nuclei as mentioned in figure legends and in Supplementary Table 3.

## ACKNOWLEDGMENTS

We thank Susan Gasser and Anais Cheblal for the SpotTracking and Excel Macro, Charles White for RAD51-GFP line, Anne Britt for the *sog1-1* mutant, Sashihiro Matstunaga for RAD54-eYFP line, Masaaki Umeda and Shiori Aki for the Cytrap line and Confocal Microscopy Platform from SLU, Uppsala, Sweden. We thank Marion Orsucci for help with the statistical analysis. We thank Lauriane Simon for comments on the manuscript. This work was supported by the Swedish Research Council (Vetenskapsrådet) grant number 2018-04101. We would like to thank COST ACTION CA16212 INDEPTH for the inputs and comments.

## AUTHOR CONTRIBUTION

AM, SR, FP and AS designed the experiments. AM and SvR performed the experiments. AM, SR, FP and AS analyzed the data. All authors contributed to writing the article and approved the submitted version. AM and SR were supported by the Swedish Research Council (Vetenskapsrådet) grant number 2018-04101. FP is supported by the “Laboratoires d’Excellences (LABEX)” TULIP (ANR-10-LABX-41)” and/or by the “École Universitaire de Recherche (EUR)” TULIP-GS (ANR-18-EURE-0019).

---

**Figure S1:**
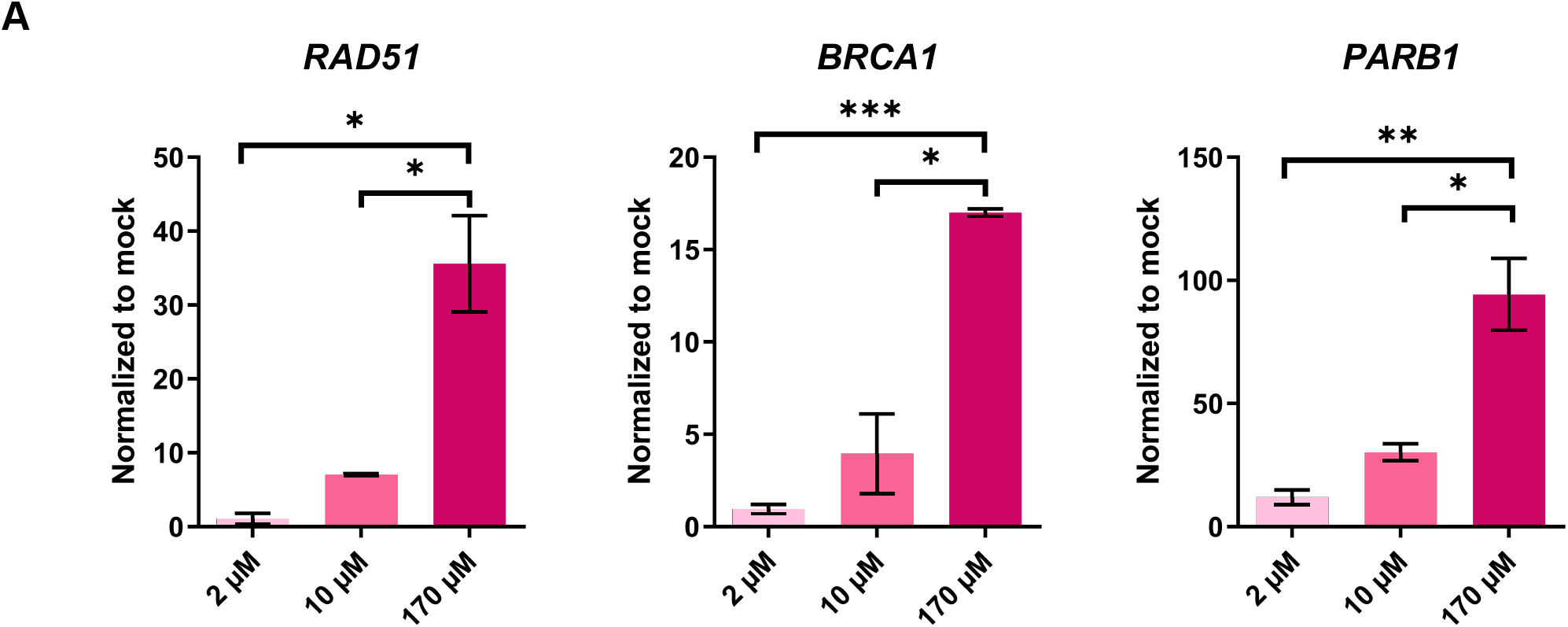
Zeocin induces expression of DDR genes in a dose-dependent manner. Expression analysis of DDR responsive genes RAD51, BRCA1 and PARB1 in 7-day-old seedlings treated with different zeocin concentrations. Two to three independent biological replicates were performed. Values represent means ± SEM. Student t-test, *P < 0.05 **P < 0.01 ***P < 0.001.

**Figure S2:**
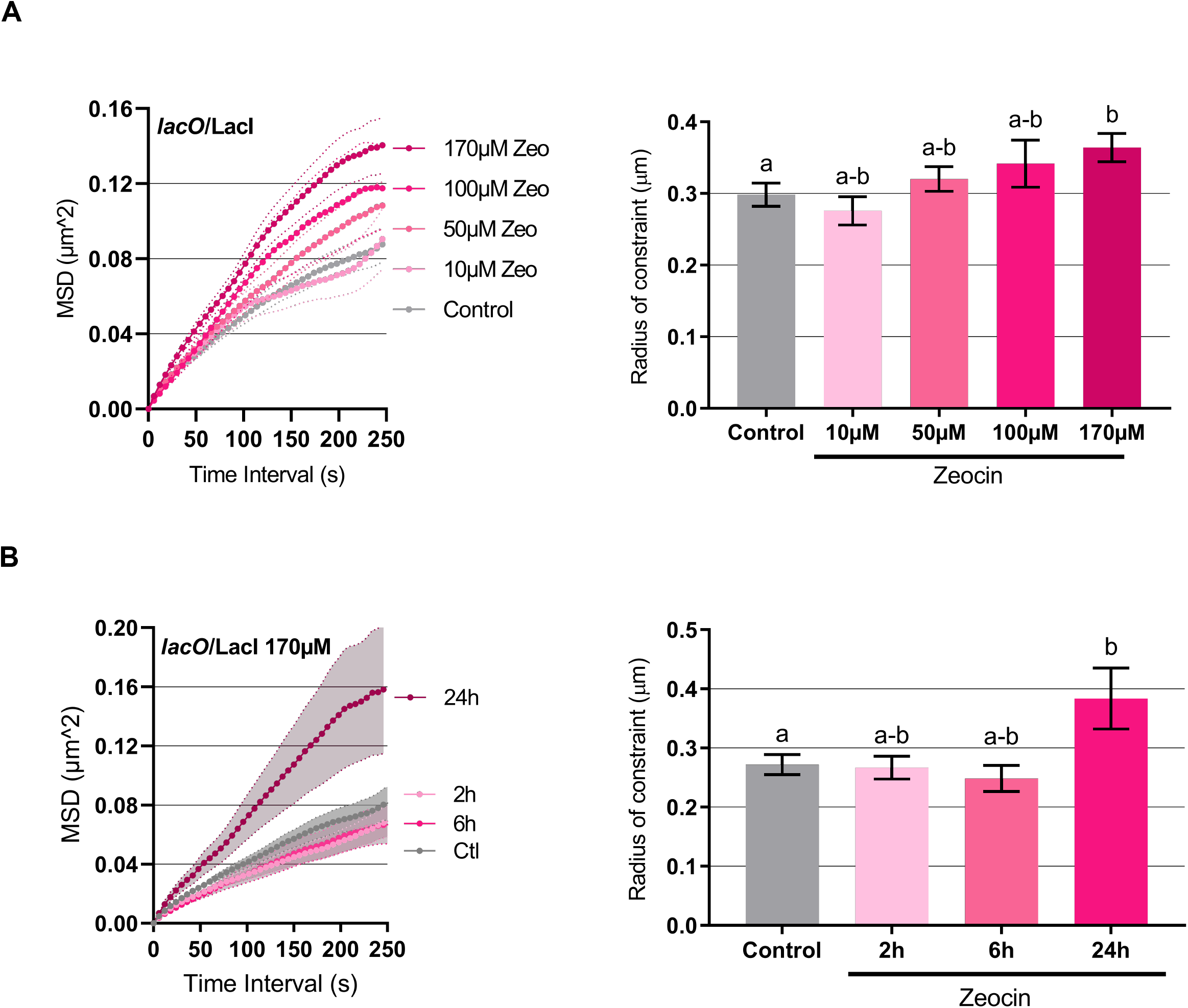
Chromatin mobility increases in treatment with a high zeocin concentration. (A-B) MSD curves and corresponding Rc histograms for: (A) *lacO*/LacI line 112 in control conditions and upon treatment with different zeocin concentrations (10μM n=97; 50μM n=111; 100μM n=29; and 170μM n=93 nuclei); (B) *lacO*/LacI line 112 in control conditions and different time zeocin treatment (2h n=39; 6h n=45; and 24h n=20); Values represent means ± SEM. Letters indicate one-way ANOVA followed by Bonferroni’s correction (p < 0.05)

**Figure S3:**
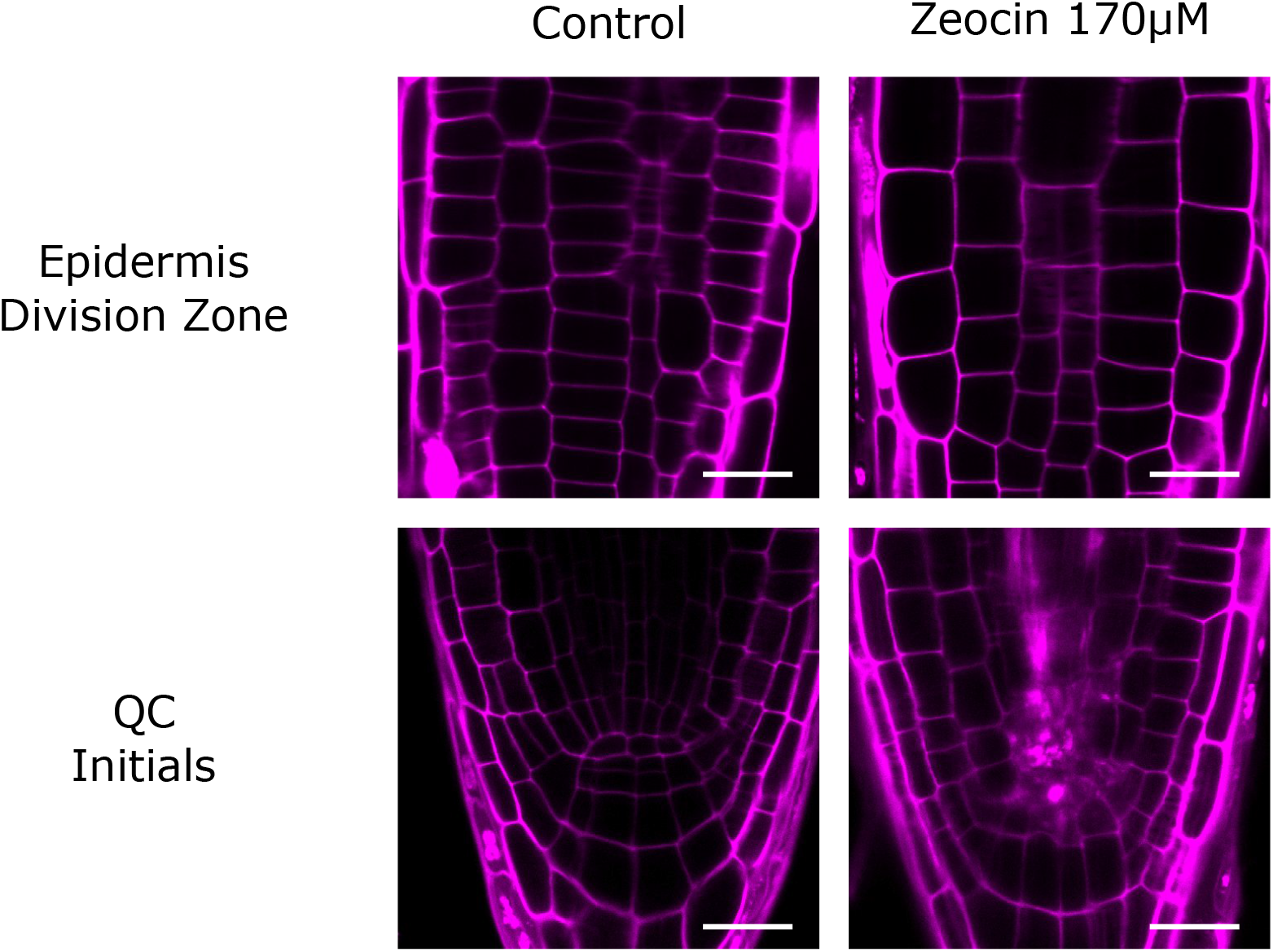
Genotoxic stress upon zeocin induces cell death in QC but not in epidermal cells from the division zone. Representative images of roots stained with PI, which marks the outline of living cells but enters dead cells, from epidermal cells in division zone and stem cell niche (QCs and Initials) from 7 days old Col-0 seedling after 24 h of zeocin treatment compared to non-treated samples (Control). Scale bar, 20 μm.

**Figure S4:**
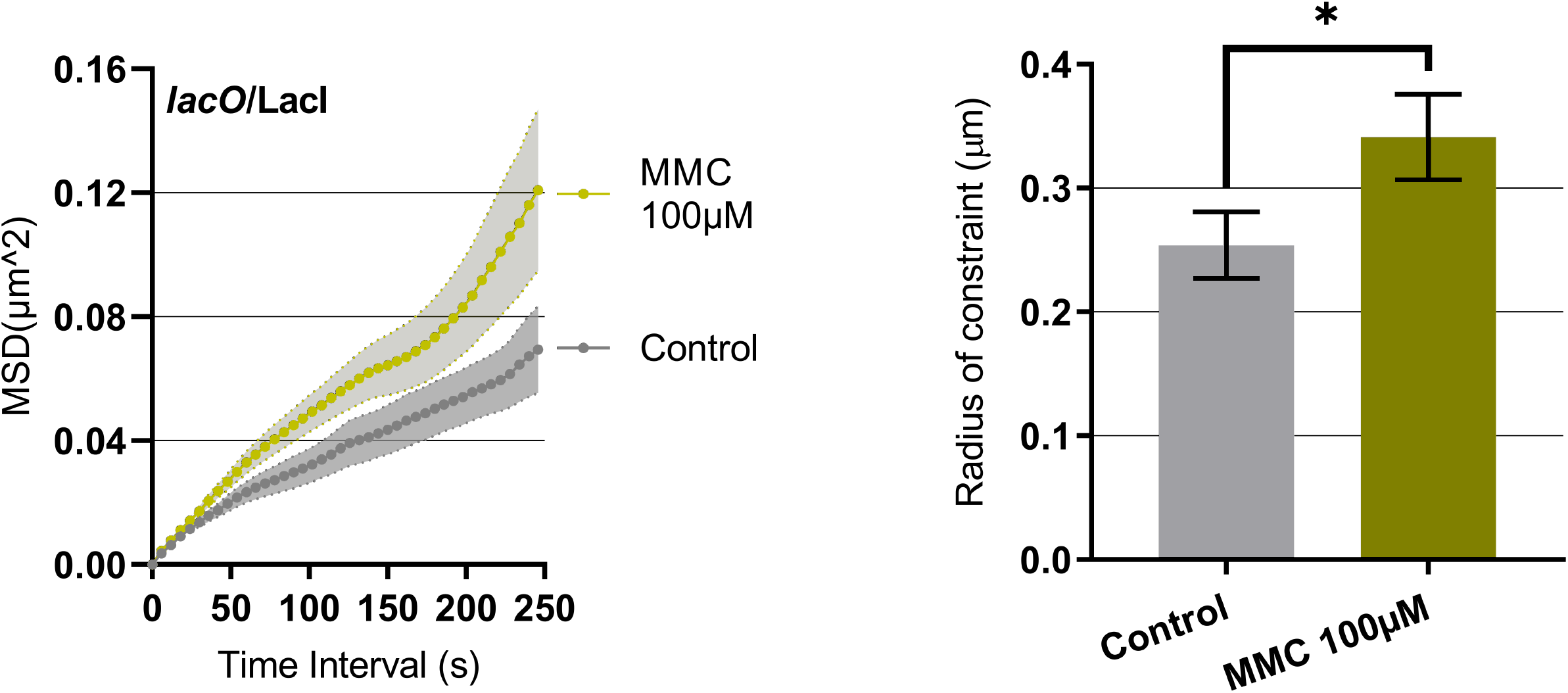
Chromatin mobility increase upon MMC treatment. MSD analysis of *lacO*/LacI lines based on time-lapse experiments of nuclei upon MMC. Radii of constraint were calculated from MSD curves. Control n=32; 100μM n=30 nuclei. Values represent means ± SEM. Student t-test, *P < 0.05.

**Figure S5:**
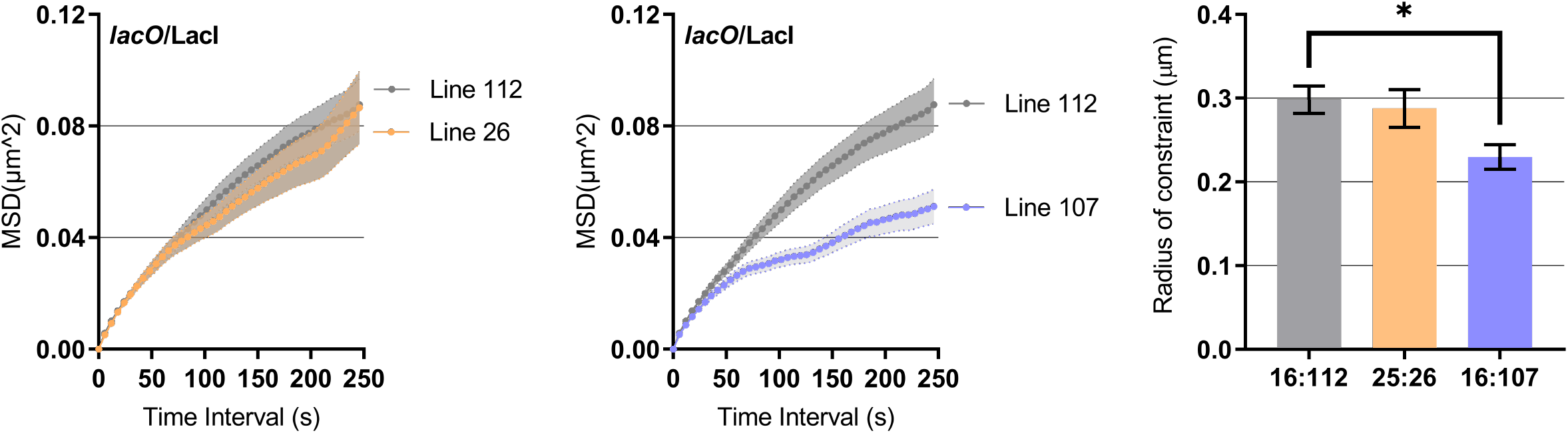
Chromatin mobility for additional *lacO*/LacI lines in control conditions. MSD analysis of *lacO*/LacI line 112 (n=116 nuclei) compared to lines 26 (n=52 nuclei) and 107 (n=53 nuclei). Radii of constraint were calculated from MSD curves. Values represent means ± SEM. Student t-test, *P < 0.05.

**Figure S6:**
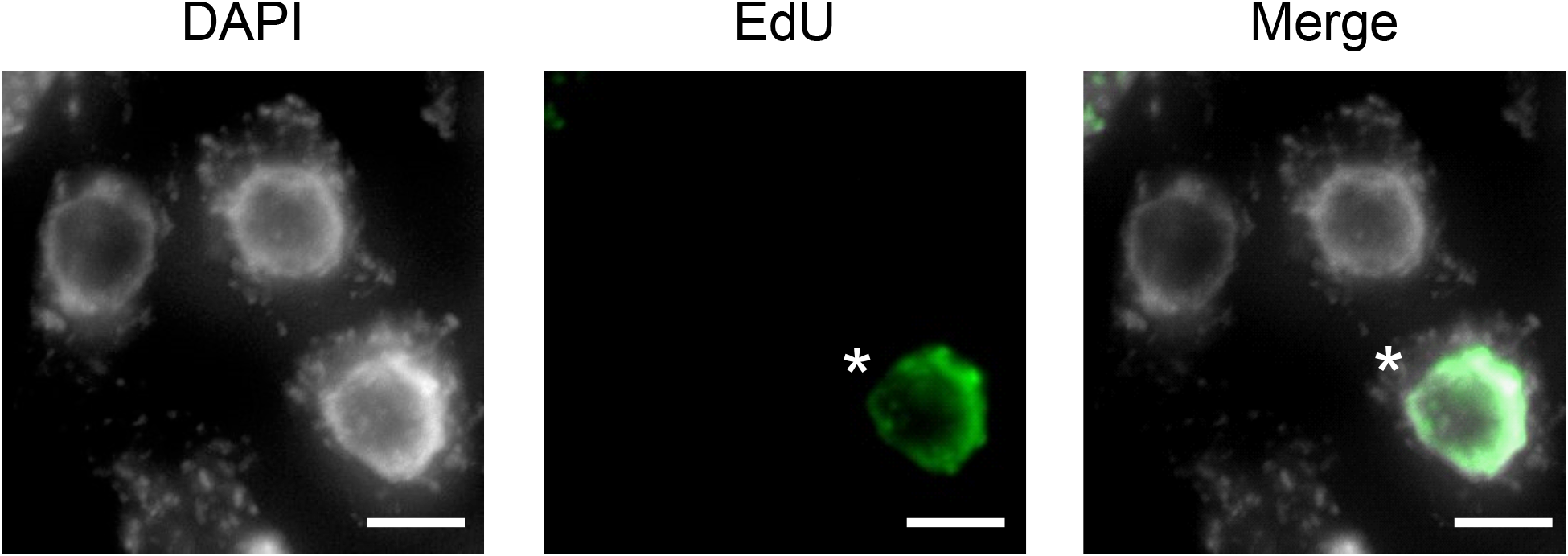
Quantitative analysis of EdU incorporation on *Arabidopsis thaliana* roots. DAPI-stained (DNA) and EdU-labelled cells from the meristem region after roots were incubated for 6h in EdU. Asterisk indicate cells showing EdU signal. Scale bar 10 μm.

**Figure S7:**
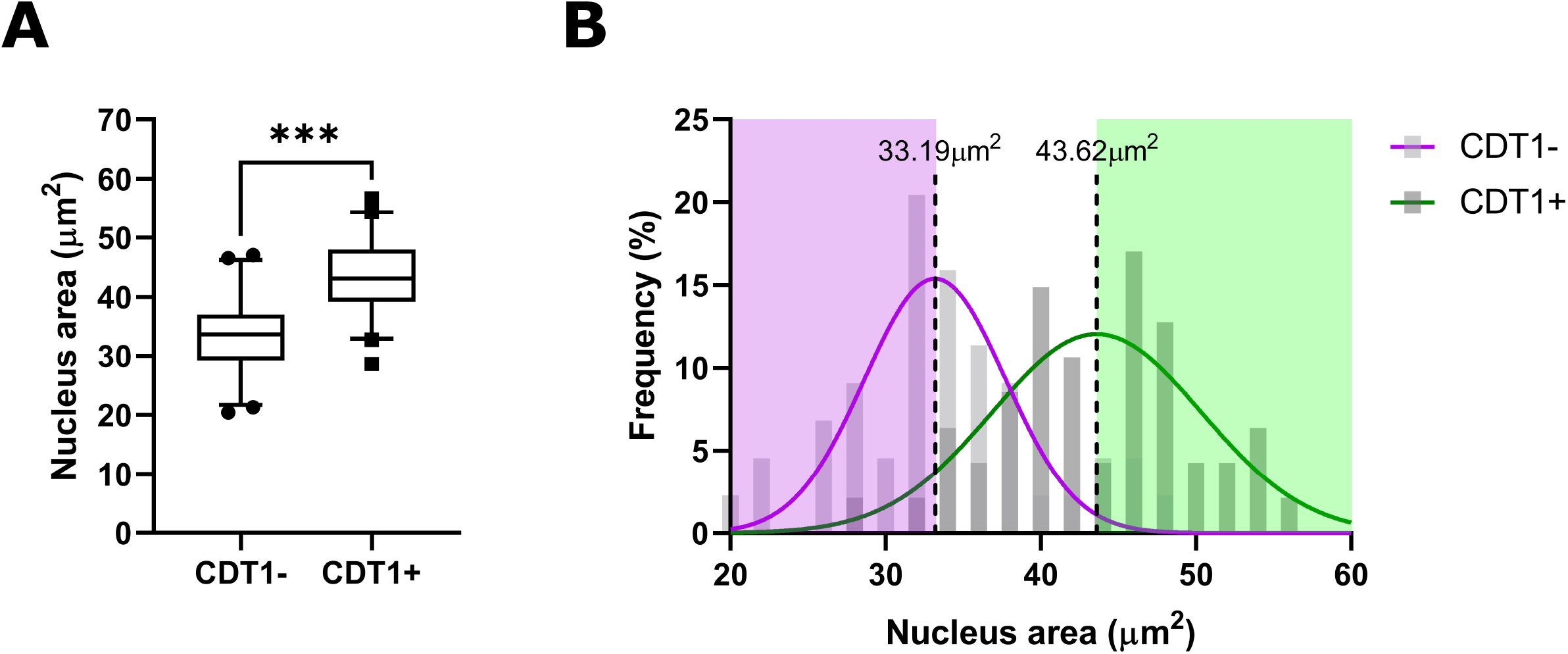
Nucleus area in S/G2 versus G1 cells. (A) Nuclear area quantification in nuclei with and without CDT1 signal. Nuclei from epidermal trichoblast root cells from *lacO*/LacI lines were measured from optical slices obtained from confocal microscopy imaging. (B) Frequency distribution in percent for the nuclear area (μm2) for nuclei with (dark gray) and without (light gray) CDT1 signal. Each curve was fitted by a Gaussian function (CDT1-in purple; CDT1+ in green). The median value of CDT1-/+ distribution has been used as thresholds to determine cells in G1 or S/G2 phase. Values represent means ± SEM. Student t-test, ***P < 0.001.

**Supplementary Table 1:**
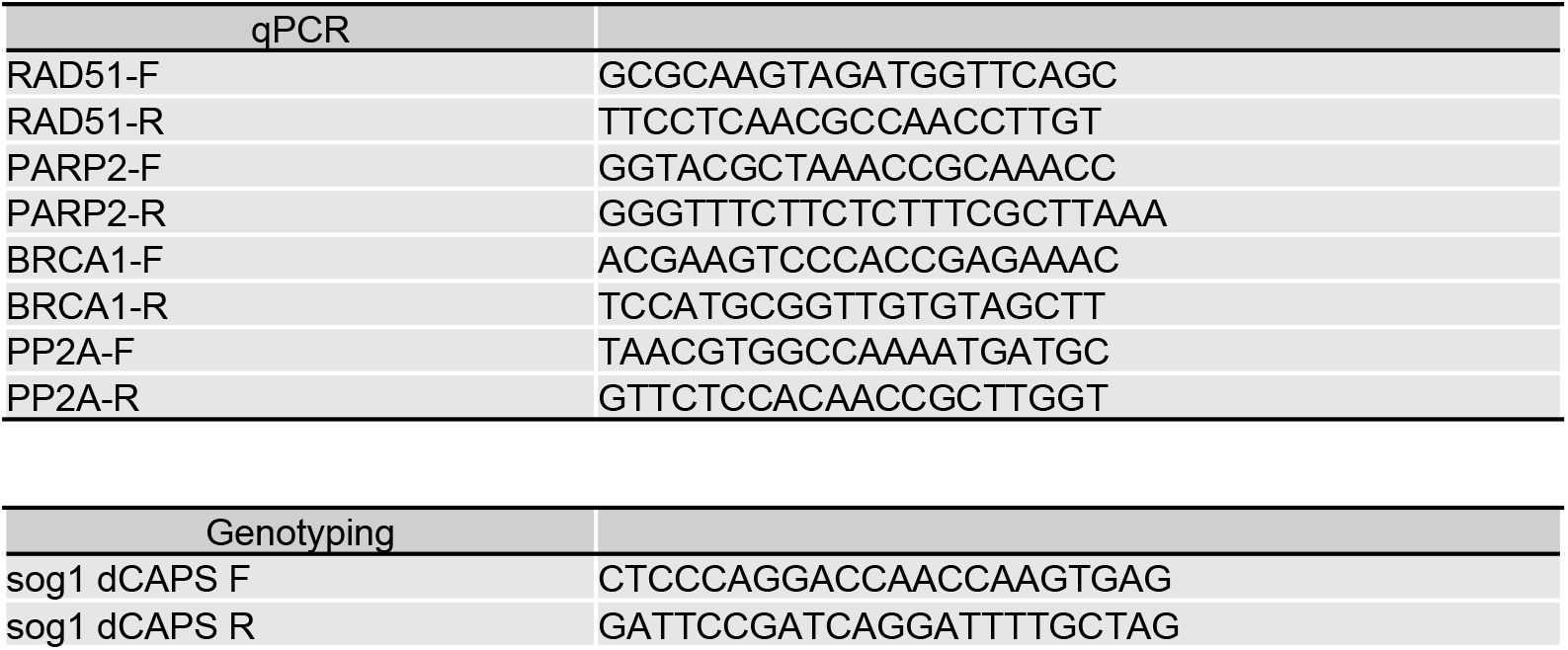
Primers Used in This Study

